# Lamellar Schwann cells in the Pacinian corpuscle potentiate vibration perception

**DOI:** 10.1101/2024.08.23.609459

**Authors:** Yuh-Tarng Chen, Dominica de Thomas Wagner, Alastair J. Loutit, Ali Nourizonoz, Mary-Claude Croisier-Coeytaux, Jérôme Blanc, Graham Knott, Kuo-Sheng Lee, Daniel Huber

## Abstract

Pacinian corpuscles are among the most sensitive mechanoreceptors found in vertebrates and they are tuned to vibrations in the highest perceptible frequency range (100-2000Hz). One of their anatomical hallmarks is the onion-like cell layers surrounding the central axon. The innermost layers consist of ∼60 densely packed lamellar Schwann cells (LSCs), whose function remains largely unknown. Using high-resolution 3D electron microscopy we found that LSCs in Pacinian corpuscles of the mouse hindlimb do not form concentric rings, but complex, multilayered and intertwining assemblies that are connected via an estimated 5805.1 desmosomes and 4142.5 gap-junctions. LSCs make multiple converging contacts with the afferent axon and its protrusions with desmosomes. Using optogenetic manipulations of LSCs we demonstrate that their activation does not only drive reliable time-locked spiking in the axon, but that their inactivation significantly elevates the thresholds in-situ and increases perceptual thresholds behaviorally. Together these findings provide evidence that LSCs are a key element of somatosensory processing, actively potentiating mechanosensitivity in Pacinian corpuscles.

**Highlights:**

- High-resolution electron microscopy reveals details of the Pacinian corpuscle
- Lamellar Schwann cells form claw-like structures with converging axonal contacts
- Schwann-cell modulation bidirectionally affects neural coding of Pacinian afferent
- Inactivation of lamellar Schwann-cells increases perceptual thresholds

## Introduction

The somatosensory system is remarkably well-adapted to extract vibratory stimuli from the environment, which arise from surrounding movement and when actively exploring surface textures and frictions. In vertebrates, vibrations are transduced into electrical impulses by different mechanoreceptor end-organs, such as Merkel cells, Ruffini, Krause, Meissner and Pacinian corpuscles, or Lanceolate endings (Handler and Ginty 2021; Qi et al. 2024). Pacinian corpuscles (PCs) are uniquely sensitive to high-frequency substrate vibrations in the range of tens of nanometers, despite being typically located near the hypodermis and periosteum and thus further away from the skin surface compared to most other receptor types (Bell, Bolanowski, and Holmes 1994; Handler and Ginty 2021). To uncover the mechanism by which PCs achieve such exquisite vibration sensitivity requires not only a detailed understanding of their cellular structures and their respective interactions, but also functional manipulations assessing their contribution to physiological activation.

PCs are extremely large compared to other mechanoreceptors (∼2mm in human), and consist of a central elliptic cylindrical axon that often ends in a bulbing and bifurcating tip (Handler et al. 2023). This afferent axon terminal expresses high densities of mechanosensitive channels (Handler et al. 2023) and is tightly wrapped with lamellar Schwann cells (LSCs), also known as non-myelinating sensory Schwann cells (Ojeda-Alonso et al., 2024) or terminal Schwann cells (Handler et al., 2023), forming the inner core. In a transverse section, the Schwann cell lamellar layers appear to form two concentric semilunar halves separated by two clefts, which span across the inner core. Protrusions extend from the axon into the cleft region (Bolanowski, Schyuler, and Slepecky 1994; Handler et al. 2023; Pease and Quilliam 1957). The inner core is surrounded by three groups of concentrically arranged layers: first the intermediate layer, then the outer core layers, and finally the capsule. The intermediate layer consists of endoneurial CD34+ cells (García-Piqueras et al, 2017), whereas the outer core and capsule layers are made of perineural squamous epithelial cells (Shanthaveerappa and Bourne 1963).

How this ensemble of layered surrounding cells participates in mediating touch sensation remains unclear. Early experiments showed that the removal of the outermost layers (decapsulation) substantially affects PC kinetics and rapid adaptation (Bell, Bolanowski, and Holmes 1994; Mendelson and Lowenstein 1964). Previously proposed roles of the LSCs include the maintenance of the ionic or metabolic microenvironment (Dubový 1986; Bell, Bolanowski, and Holmes 1994), the mediation of rapid adaptation of afferent firing (Pawson, Pack, and Bolanowski 2007; Pawson et al. 2009) and, most recently, active agents in initiating pain and somatosensation (Ojeda-Alonso et al. 2024; Abdo et al. 2019). The innermost LSC layers have been suggested to play a central role in mechanotransduction in Meissner corpuscles (Iwanaga et al. 1982; Landcastle et al. 1991; Nikolaev et al. 2023; Abdo et al. 2019; Handler et al. 2023; Bell, Bolanowski, and Holmes 1994). In the avian Meissner corpuscle homolog, it has been shown that LSCs are mechanosensitive and excitable (Ziolkowski, Gracheva, and Bagriantsev 2022; Nikolaev et al. 2023, 2020). Also, recent work demonstrated that LSCs surrounding mechanoreceptor nerve-endings within the Meissner’s corpuscle and in hair follicle lanceolate endings set perceptual thresholds for touch (Ojeda-Alonso et al. 2024). Specific subcellular elements such as tethers (Ojeda-Alonso et al. 2024; Hu et al. 2010) that assist in opening mechanically gated channels such as Piezo2 or adherens junctions that help to stretch cellular membranes have also been proposed to play roles in mechanosensation, particularly in rapidly adapting mechanoreceptors (Handler et al. 2023).

Taken together, LSCs are ideally situated to potentiate mechanotransduction, but it remains unknown if they participate in the PC’s exquisite sensitivity to high-frequency vibration. Here, we combine high resolution volumetric electron microscopy and optogenetic manipulation to directly assess the contribution of LSCs to the transduction of mechanical stimuli.

## Results

### Ultrastructure of the Pacinian corpuscle and the lamellar Schwann cells

To understand the detailed morphology of Pacinian LSCs, we conducted an ultrastructural analysis using serial block-face scanning electron microscopy (SBF-SEM). PCs were dissected from the hindlimb periosteum of a mouse (**Fig. S1**), processed for electron microscopy (see **Methods**). We first imaged a complete PC at “lower” resolution (voxel size: 40 x 40 x 200 nm, **Fig. 1A**) and manually reconstructed the different primary components (**Fig. S2**). This three-dimensional reconstruction revealed what had previously been extrapolated from single section transmission electron microscopy (TEM) and identified by conventional light microscopy (Pacini 1840; Pease and Quilliam 1957; Bolanowski, Schyuler, and Slepecky 1994; Spencer and Schaumburg 1973; Bell, Bolanowski, and Holmes 1994), that a PC consists of tightly packed LSCs wrapping around an axon, which in turn are surrounded by the outer core consisting of multiple concentric layers of flattened cells, which are subdivided into an intermediate layer, an outer core and a capsule (**Fig. 1A**). While the axon is a straight elliptic cylinder in the proximal portion of the PC, distinct bulbings and bifurcations are observed in the distal portion (ultraterminal) of the PC (**Fig. S2A**).

**Figure 1.**
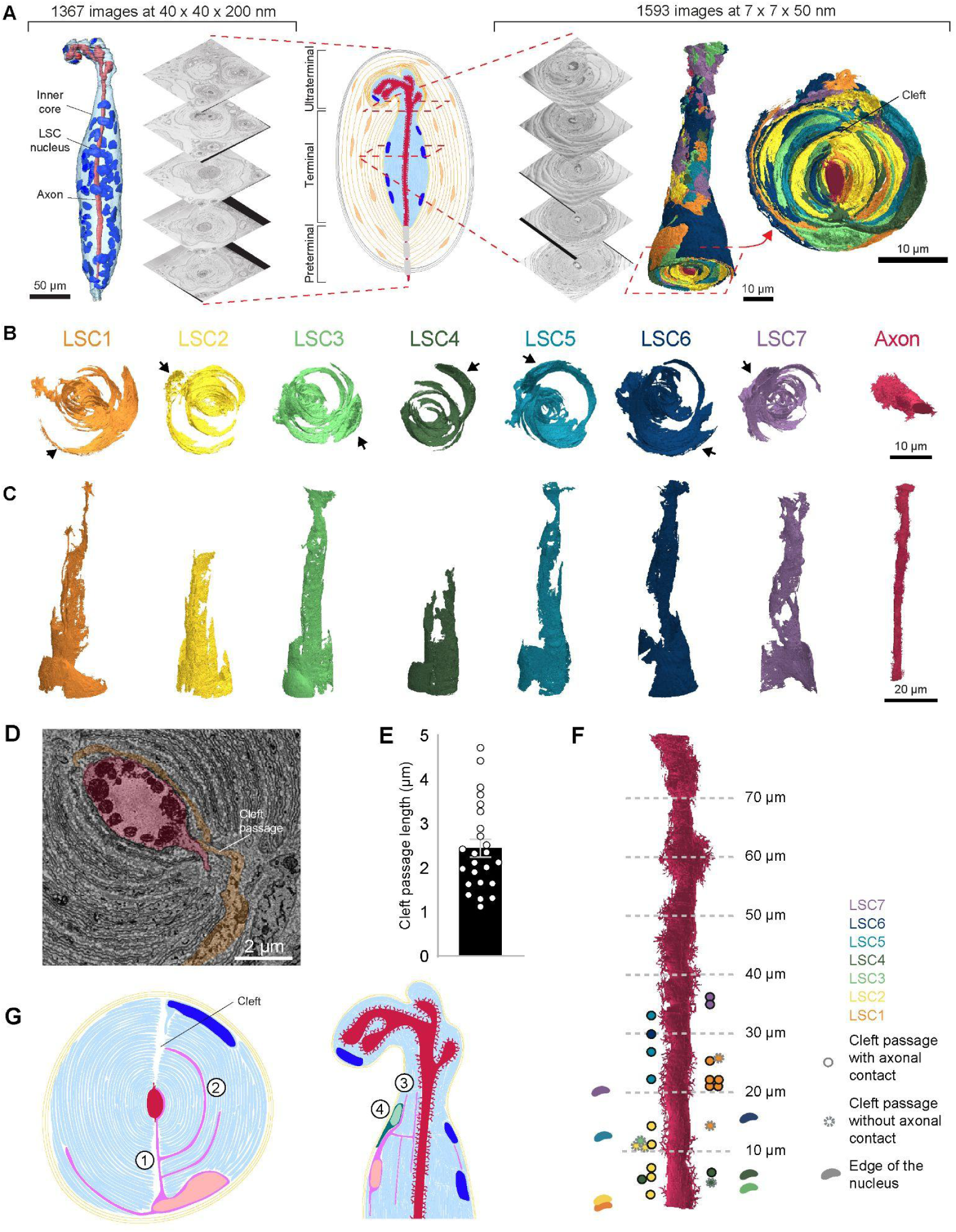
High resolution SBF-SEM reconstructions reveal the complex morphology and surface area of the lamellar Schwann cells comprising the PC inner core. **(A)** Schematic showing the two image stacks acquired with SBF-SEM. The first allowed for reconstruction of all components of the PC (**Fig. S1**). Here we display the inner core region containing 63 LSC nuclei, with the axon in the middle. The second SBF-SEM stack is a 75μm subsection of the inner core, which was acquired at higher resolution and allowed for the reconstruction of individual LSCs. The renderings of all seven reconstructed LSCs together shows the interleaving of the layers from different cells. **(B-C)** Three-dimensional renderings of the seven individual LSCs and axon reconstructed from the higher resolution stack, seen from an x-y (B) and x-z (C) perspective. The black arrows show the position of the individual LSC nucleus. **(D)** Example cleft passage from LSC1. The cleft passage makes a contact point with an axon protrusion and leads to the formation of a new layer closer to the axon. **(E)** Length of LSC segments in the cleft. **(F)** Schematic showing LSC nuclei positions and cleft passages, including their locations and whether each cleft passage has direct axon contact. **(G)** Schematic displaying structural characteristics of LSCs. 1. LSCs have branches that directly approach the axon from the periphery of the inner core by cutting through the cleft. 2. LSCs form many layers in the inner core. 3. LSC layers extend proximally and distally within the inner core. 4. Those distal LSCs (located closer to the ultraterminal) are often positioned relatively inwardly compared to the proximal LSCs.

Although at this resolution we were able to reconstruct the overall shape of the inner core, and identify the location and number of LSC nuclei (63, **Fig. 1A**), it was not possible to distinguish individual LSC membranes. We therefore sought to reconstruct a more detailed morphology of individual LSCs, to determine how they interact with the axon and with other LSCs. For this purpose, we processed and imaged a subsection of a PC at higher resolution (voxel: 7 x 7 x 50 nm; **Fig. 1A**) using SBF-SEM. The total size of the imaged block was 29 x 31 x 80 µm, which comprised the full transverse cross-section of the inner core, but not its full length. We began by manually reconstructing the axon (**Fig. S2A**) and one LSC (**Fig. S3A**) by visual annotation, but subsequently developed an image analysis and segmentation pipeline (see **Methods**) using Webknossos (Boergens et al. 2017), facilitating the automated reconstruction of seven LSCs (**Fig. S3**). When viewed in the transverse plane (**Fig. 1B**), 3D renderings of the reconstructions reveal cells that consist of multiple lamellae that wrap around the axon in a claw-like manner, with an unevenly distributed surface area (**Fig. S4A**), in stark contrast to previous descriptions of evenly distributed layers split by two clefts. The elongation of the cells along the axon is demonstrated when viewed longitudinally (**Fig. 1C**). Viewing all seven LSCs jointly (**Fig. 1A**) highlights the onion-like appearance and the manner in which lamellae from different intricately interlaced LSCs. The different claw segments converge towards the central cleft. Each LSC extends a long segment directly through the cleft towards the axonal protrusions, from which other segments extend perpendicularly, forming the lamellar layers wrapping around toward the opposing side of the axon (**Fig. 1D, G**). In total, the 7 LSCs make 24 passages longer than 1 μm in the cleft (**Fig. 1E**). Of these, 18 lead to direct contact with the axon, while the other 6 do not. This suggests that LSCs use the clefts as “shortcuts’’ to interact with the axon. We also found that most LSCs (6 of the 7 reconstructed) enter only on one side of the cleft , and this can be either on the same side as the nucleus or on the opposite side (**Fig. 1F**).

Quantification of the surface area of each LSC revealed a pattern of significantly decreasing surface area from proximal to distal regions (F(7,48) = 11.88, *P < 0.0001, n = 7, one-way ANOVA), while the axon surface area conversely increased (**Fig. S4B**). Taken together, we conclude that LSCs form stereotypical funnel-shaped, intertwined structures which not only extend in a claw-like fashion, but use the cleft as the main passage towards the axon.

### Contacts between lamellar Schwann cells and the afferent axon

The close proximity and tight claw-like wrapping of inner core LSCs suggests they may have a direct role in potentiating mechanotransduction in PCs. Yet, whether inner core LSCs directly interact with the axon, and the types of junctions that might facilitate communication between them, remains largely unknown. Here, our 3D reconstructions reveal direct contact points between LSCs and the axon and provide detailed insights into the spatial and quantitative aspects of these interactions. **Figure 2A** depicts all the contact points (defined as a distance of 30 nm or less between axon and LSC, see **Methods**) made on both the afferent axon body and axonal protrusions (**Figure 2B**). Along the axon, protrusions do not individually emerge directly from the axon, instead they are grouped together and sprout out of common trunks (**Figure 2C)**. The distribution of LSC contact points have their highest density near the middle of our imaged volume and decrease sharply near the ultraterminal (**Fig. S4C**). LSC-axon contacts are highly clustered, with each LSC having a preferred region where the axon body is contacted directly **(Fig. 2E & F)** or on the trunks and protrusions **(Fig. 2G & H)**. Overall, LSCs have greater contact surface area with the axon body, but more numerous contact points on the axon protrusions (**Fig.S4D; Table S1**). In addition, we observe that the position of the LSC nucleus determines its first contact with the axon (**Fig. 2I**). Although, the amount of LSC-axon protrusion contacts is regardless of the nucleus position relative to the cleft **(Fig. 2J)**, we found a consistent number of LSC-axon protrusion contacts across the trunks (on average 6-7 contacts) on a local scale **(Fig. 2K)**. Yet surprisingly, over half of these LSC-axon protrusion contacts were formed by only one or two individual LSCs **(Fig. 2L**). This means that contact points are not only regionally clustered but extremely selective, and comprise very few LSCs.

**Figure 2.**
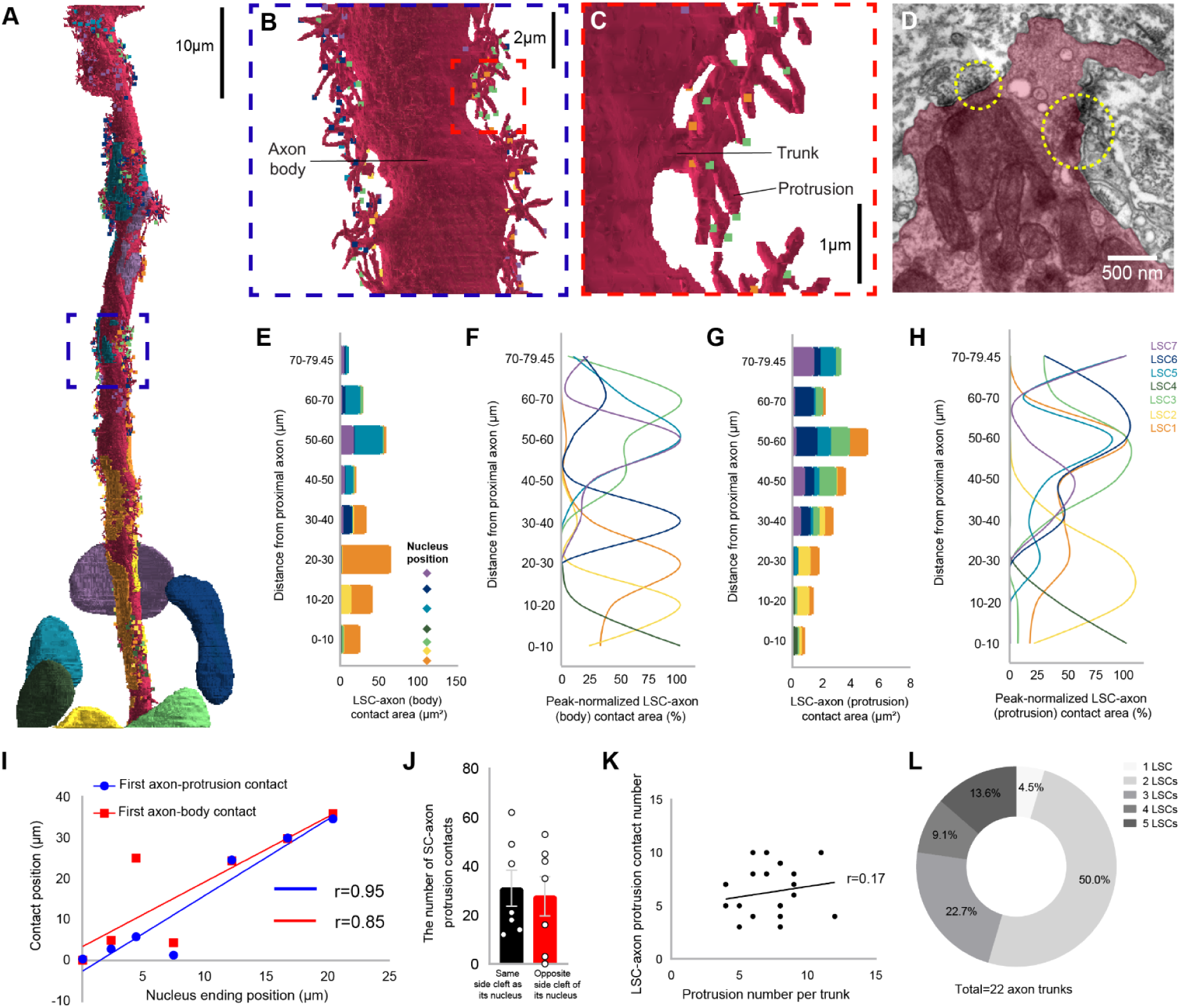
Lamellar Schwann cells form contact points with the axonal body and protrusions. **(A)** 3D rendering of SBF-SEM reconstructions, showing the position of LSC nuclei, LSC-axon protrusion contacts (color squares), and LSC-axon body contacts (color spots). **(B)** Close up of a smaller section of the axon. **(C)** Trunks that sprout into multiple protrusions line the length of the axon. **(D)** TEM section, showing electron-dense events where the LSCs contact the axon. **(E)** Accumulating distribution (stacked bar chart) of LSC-axon contact areas on the axonal body along the length of the axon. **(F)** Peak normalized distribution of LSC-axon contact areas on the axonal body along the length of the axon. **(G)** Accumulating distribution (stacked bar chart) of LSC-axon contact areas on the axonal protrusions along the length of the axon. **(H)** Peak normalized distribution of LSC-axon contact areas on the axonal protrusions along the length of the axon. **(I)** Relationship between nucleus location and first contact point between LSC and axon. With Pearson’s correlation coefficients (r) both showing significant differences (P < 0.05). **(J)** The number of LSC-axon protrusion contacts, separated by LSC nucleus location. **(K)** The number of contact points made on a trunk is independent of the number of protrusions that trunk has. **(L)** The number of LSCs making contact points on a single trunk.

Next, we asked if these contact points contain any specific structural or functional elements. For this purpose we used a cryo-embedding approach which better preserves the membranes allowing different complexes to be more easily identified in TEM. We found electron-dense structures within LSC-axon contact zones (<30 nm apart) (**Fig. 2D**), similar to those proposed to be desmosomes (Sakada and Sasaki 1984) or adherens junctions (Handler et al. 2023). However, we could not find any evidence for gap junctions or synaptic and vesicular structures between LSCs and the axon. Taken together, these data suggest that LSCs are structurally tightly linked to the axon, with no obvious elements for classical functional coupling.

### Contacts between individual lamellar Schwann cells

Given the tightly packed nature of LSCs within the inner core, we next investigated the potential connections between individual LSCs. Previous ultrastructural studies reported the presence of tight junctions (Sakada and Sasaki 1984) and gap junctions between LSCs (Rico et al. 1996; Ide and Hayashi 1987), suggesting that LSCs may reciprocally influence the electrical properties of adjacent LSCs. However, neither the density nor the distribution of either element is known. By using the above described cryo-embedding approach, we were able to clearly identify a high density of both desmosomes as well as gap-junctions between LSCs (**Fig. 3A**). These are not visible in the images taken with SBF-SEM (due to the embedding approach as well as the lower imaging resolution) we quantify instances of close apposition between LSC membranes as a means of quantifying junctions, an example of which is shown in **Fig. 3B-D**. To estimate the number of gap junctions and desmosomes within the inner core of the PC, we took a stereological approach and counted the number of these identified structures across serial, high resolution images (2nm/pixel) that covered an area of 680 µm^2^. We found the density of gap junctions to be 29.4 /1000 µm^3^ and 41.2 /1000 µm^3^ for desmosomes. We extrapolated these densities to the entire inner core volume of PC1, 140900.72 µm^3^, and therefore estimate a total of 4142.5 gap junctions and 5805.1 desmosomes connecting the approximately 60 LSCs. This would mean that each LSC has on average 69.0 gap junction and 96.8 desmosome contacts to other LSCs. Taken together, these results suggest that the densely intertwined LSCs not only form a mechanical, but also functionally tightly coupled network.

**Figure 3.**
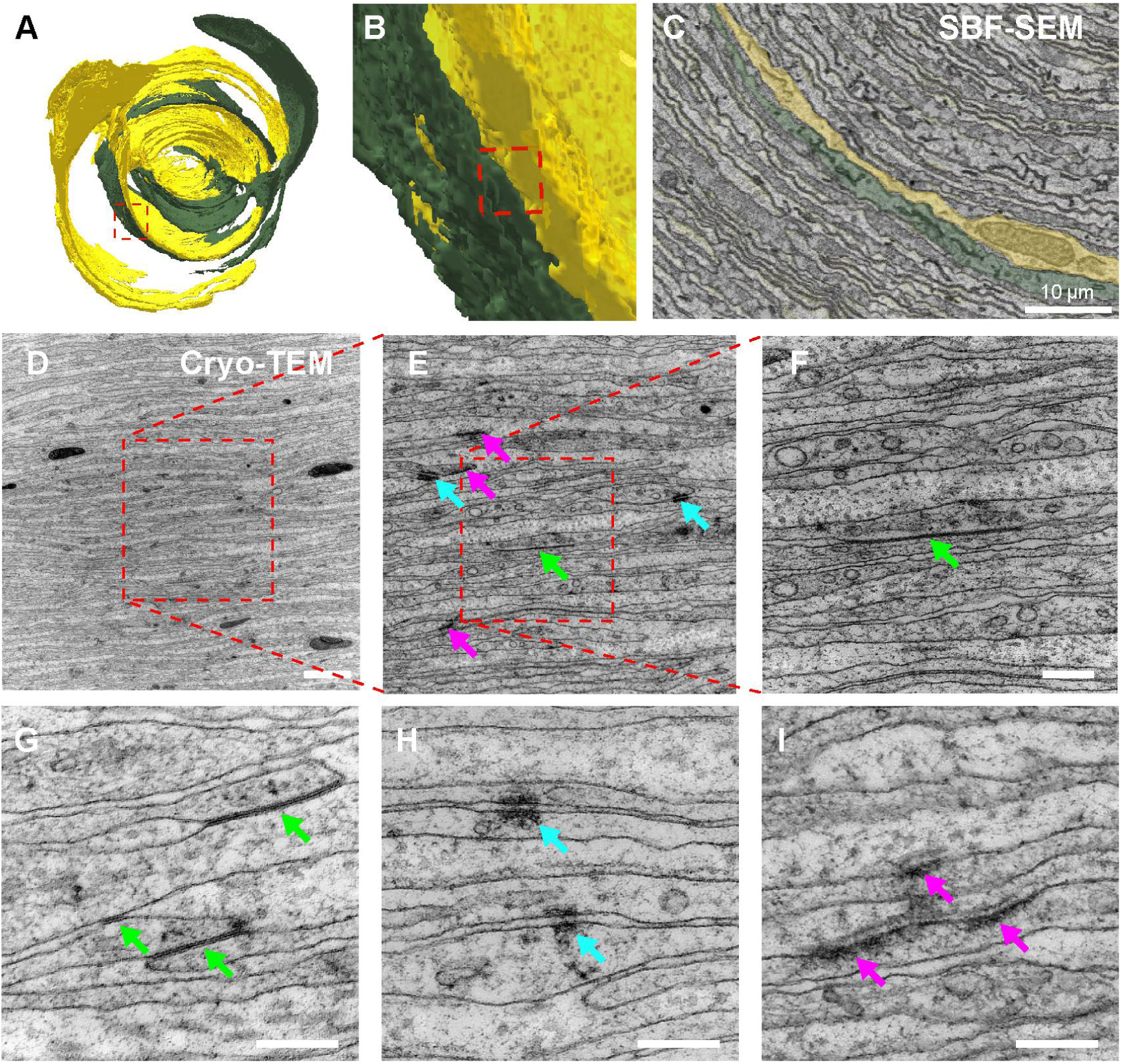
LSCs make gap junctions with other LSCs. **(A)** Three-dimensional renderings of two LSC (LSC2, LSC4), the red box denotes the locations of an LSC-LSC contact. **(B)** Close-up of the site of the LSC-LSC contact in (A). **(C)** SBF-SEM image showing the contact between LSC2 and LSC4. **(D-F)** Cryo temperature staining TEM of increasing magnification indicating the appearance of electron dense structures including gap-junctions (green arrows), desmosomes (cyan arrows) and hemidesmosomes (pink arrows). **(G)** Gap-junction (green arrows) are defined by symmetrical close appositions of thin electron dense structures with no space between the membranes. **(H)** Desmosomes (cyan arrows) are marked by short symmetrical electron dense element with a visible gap. **(I)** Hemidesmosomes (pink arrows) are electron dense structures only found asymmetrical on a single membrane oriented towards the extracellular space. .

### Contact surface area is prioritized between lamellar Schwann cells but not with the axon

The density of gap junctions between LSCs, but lack of gap junctions between LSCs and the axon, led us to ask the question if gap junction and desmosome densities are related to the total available contact surface area between LSC-LSC and LSC-axon contacts. We found that the number but not the area of LSC-axon protrusion contacts **(Fig. S5A, B),** and the LSC-LSC contact area (**Fig. S5C**) appears to depend on the size of the LSC surface area (**Fig. S5D**). Furthermore, we observed that most LSC-LSC contacts have similar contact surface area over the proximal to distal axis **(Fig. S6A, B),** but some individual LSCs have a greater overall LSC-LSC surface area (such as LSC6). Taken together, LSC-LSC contacts appear to maximize the total surface area between them, which may increase the probability for gap junction connections and thus rapid and high fidelity LSC-LSC communication. In contrast, LSC-axon connections appear to maximize the number of contacts onto the axonal protrusions, and these contacts prioritize mechanical rather than functional coupling evidenced by high densities of desmosomes and an absence of gap junctions.

### Lamellar Schwann cells contribute to the vibration coding

To probe the functional impact of this connected network of LSCs on the axonal activity in PCs, we generated mice expressing channelrhodopsin (ChR2) or Archaerhodopsin-3 (ArchT) in ETV-1 positive LSCs to excite or inhibit these cells with light (**Fig. 4A-B**). We first used blue light to selectively activate LSCs in the PCs on the fibula whilst making single-unit recordings from PC-innervating Aβ-fibers of the sciatic nerve, known as Pacinian corpuscle low threshold mechanoreceptors (PC-LTMRs). In ETV-ChR2 mice, blue light reliably evoked ultra-short latency spikes in nearly 90% of the PC-LTMR (14/16), and other afferent types, including high-threshold mechanoreceptors, rapidly-adapting receptors and hair follicles, if the light was directed to their receptive fields (**Fig. S7)**. Light stimulation of LSCs evoked intensity-dependent spiking of the PC-LTMR (**Fig. 4C-D**). PC-LTMRs recorded from ETV-ChR2 mice respond faster to light stimulation than to vibration with a piezo actuator (**Fig. S7C**). This broad range of first spike latencies to blue light stimulation (maximum up to 40 ms) was never observed in case of mechanical stimulation. The spike latencies (as a function of light intensity) in Schwann cells appear to be very different from those in cortical neurons, which are normally much shorter (5 ms, (Huber et al. 2008)), suggesting a complex mechanism in between the activation of the LSCs and the subsequent activation of the axonal afferent. On the other hand, the extremely short minimal latencies for the intense light activation suggest a tight physiological coupling between ETV+ LSCs and the PC receptor ending.

**Figure 4.**
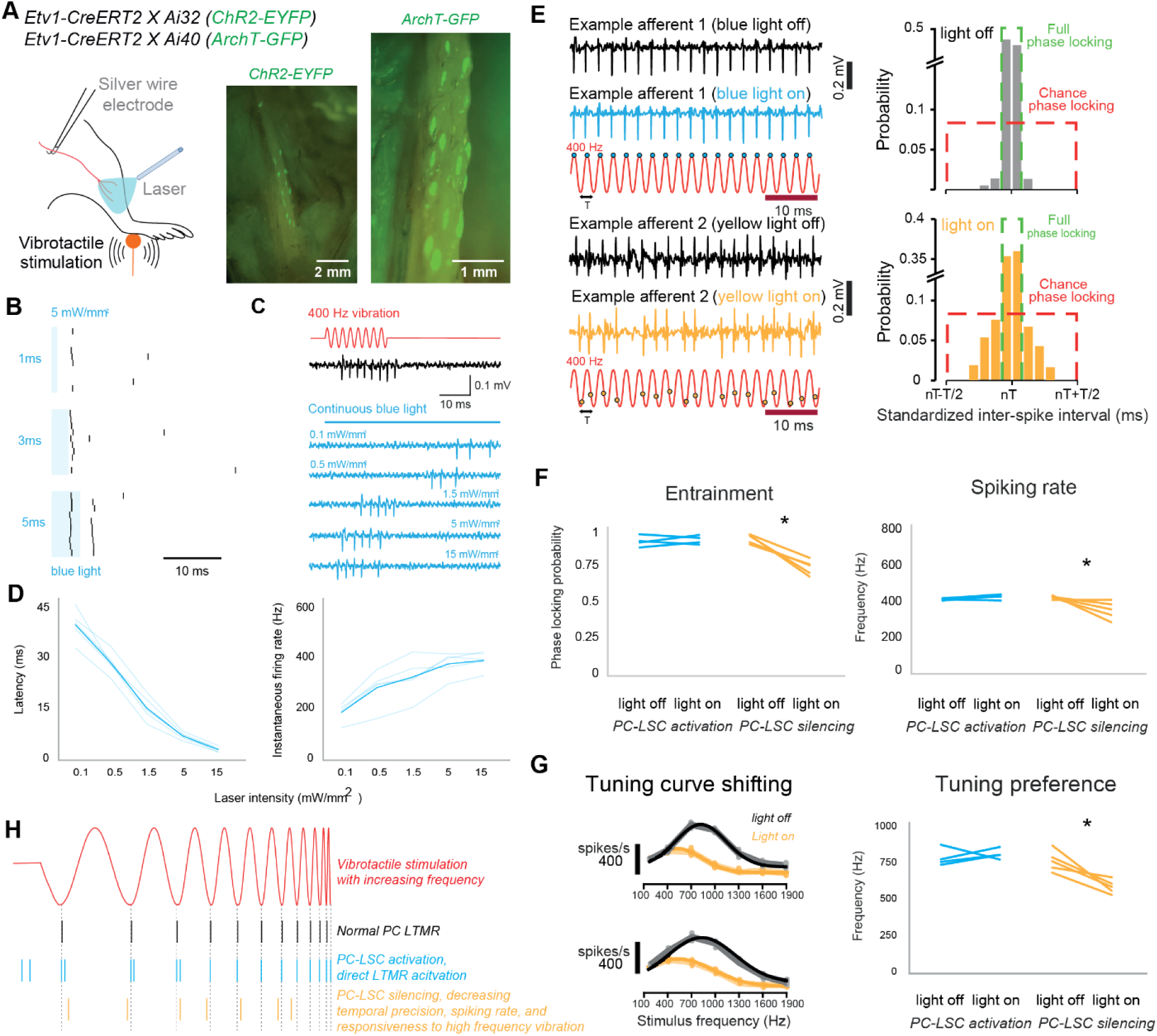
Optogenetic excitation and inhibition of lamellar Schwann cells in PC. **(A)** Schematic diagram of manipulating Schwann cells in the exposed PC during afferent recording. PCs on the fibula in a mouse hindlimb (each bright green spot is one PC), identified by EYFP or GFP expression in the inner core structures (green). **(B)** With blue laser power at 5 mW/mm^2^, the latency of light-driven spiking of the afferent is very consistent regardless of stimulus pulse duration. **(C)** Example of spiking from PCs stimulated by blue light versus a mechanical stimulus. **(D)** Intensity-dependent activation from five recorded afferents. Left, first spike latencies for PC afferent from optogenetic activation of Schwann cells. Right, instantaneous firing rate during the optogenetic activation of Schwann cells. **(E)** Left, example recording of two afferents (Top, ChR2+; Bottom, ArchT+) in response to 400 Hz vibration at the hindlimb, with light off (black) or with light on (blue/yellow). The timing of its action potentials (blue/yellow dots) relative to periodic cycles of the 400-Hz vibration (red) shows the degree of entrainment. Right, with yellow light stimulation, the distribution of standardized inter-spike intervals of an example afferent at 400 Hz vibration show decreasing degree of phase-locking probability. **(F)** Left, the phase-locking probability of afferents affected by optogenetic activation or inactivation of PC-LSCs (paired-sample t-test, *P < 0.0001). Right, the spiking rate of afferents affected by optogenetic activation or inactivation of PC-LSCs (paired-sample t-test, *P < 0.0001) in ETV-ChR2 (n = 4) and ETV-ArchT mice (n = 5). **(G)** Left, tuning curve shifting of 2 example afferents after inactivation of PC-LSCs by yellow light. Right. frequency tuning of afferents affected by optogenetic activation or inactivation of PC-LSCs (paired-sample t-test, *P < 0.001). **(H)** Schematic model showing how PC-LSCs contribute to the coding of vibration. The additional spikes in the case of PC-LSC activation (blue) illustrate the enhanced excitability of the PC-LTMR.

To evaluate the contribution of lamellar Schwann cells to mechanosensitivity in PC-LTMRs, we compared the activation by vibrotactile mechanical stimuli in the same neuron with or without light-evoked activity (**Fig. 4E-G**). With bi-directional optogenetic manipulation of LSCs in Pacinian corpuscles, we discovered the temporal precision spiking, spiking rate, and the tuning preference were all decreased in the case of LSC-inhibition. With the decreased activity in LSCs, PC-LTMR did not only drop its precise temporal code for vibration, but also shifted their peak response to the lower frequency compared to the normal PC-LTMR (Lee et al. 2024), suggesting that the membrane potential of LSCs in the Pacinian corpuscle are critical for high frequency vibration sensing (**Fig. 4E-G**). Taken together, our results show that LSCs play a critical role in neural coding of Pacinian corpuscles (**Fig. 4H**).

### Lamellar Schwann cells contribute to ultrasensitivity of perceptual thresholds for vibration

We next used light-induced activation and silencing to ask if ETV+ LSCs within PCs contribute to the ultrasensitivity of vibrotactile stimuli. In ETV-ChR2 and ETV-ArchT mice we evaluated the sensitivity using a 400 Hz sinusoidal stimulus with a linearly increasing amplitude (**Fig. 5A**). We measured the amplitude of vibration for the first spike before and after 10 minutes of continuous blue and yellow light was focused on the corpuscles. This method was used in the previous studies to demonstrate the physiological impact of Schwann cells on the sensitivities of various mechanoreceptors (Ojeda-Alonso et al. 2024; Abdo et al. 2019). In ETV-ArchT mice, all of the PC-LTMRs showed a significant elevation in mean mechanical threshold during light stimulation compared to baseline (**Fig. 5B**). Our data thus supports the hypothesis that ETV+ LSC activation within PCs lowers the threshold and increases vibration sensitivity of the corpuscles.

**Figure 5.**
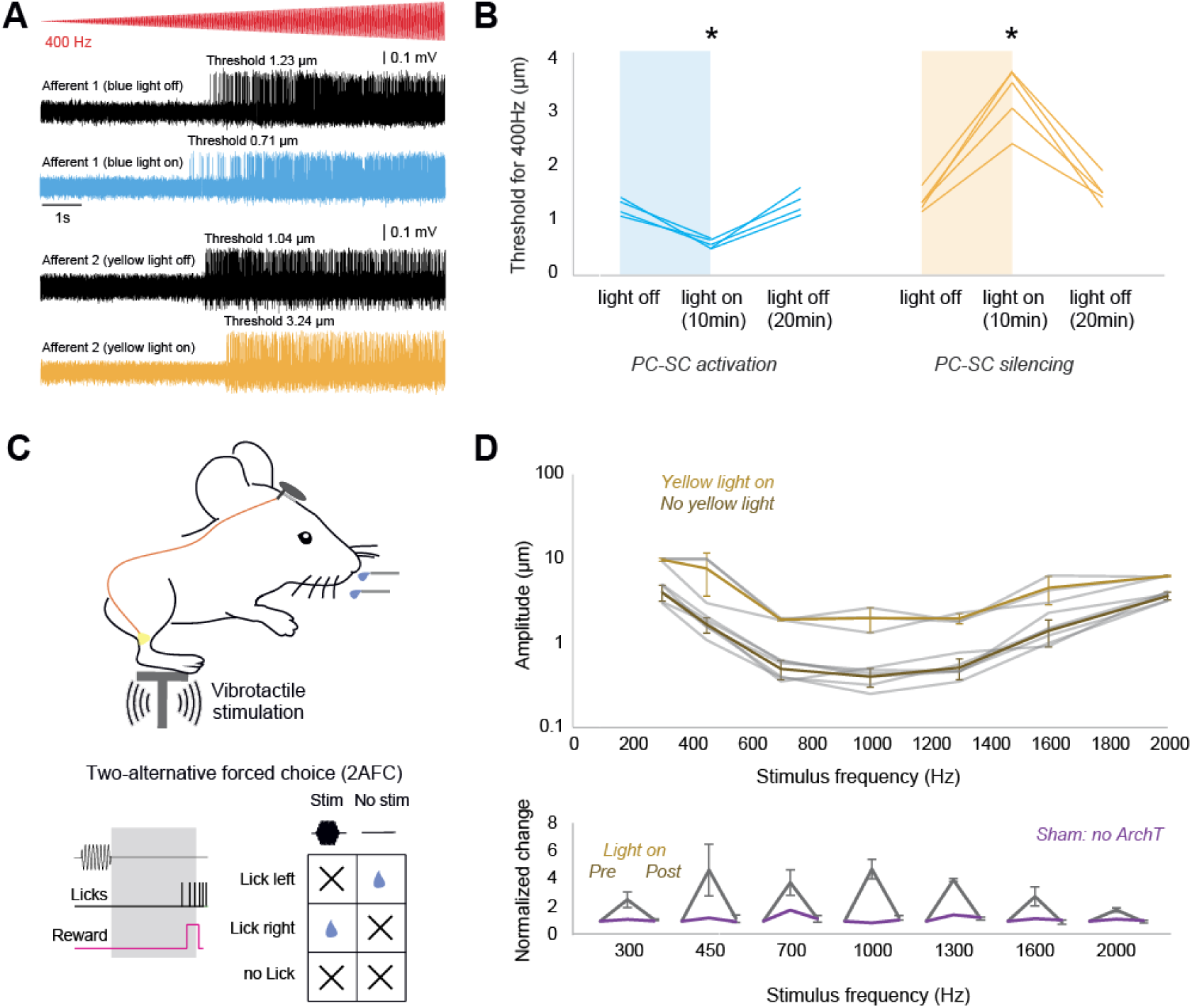
Schwann cells determine the perceptual threshold for vibration. **(A)** Mechanoreceptor spiking rates in response to 400 Hz vibration stimulus before and after optogenetic activation or inhibition of Schwann cells. **(B)** Mechanical threshold for first spike for afferents recorded in ETV-ChR2 (n = 4) and ETV-ArchT mice (n = 5). An decrease in the force necessary to evoke the first action potential was observed in afferents recorded from ETV-ChR2 mice, but the effect was revered in ETV-ArchT mice. One-way ANOVA, *P < 0.05, Bonferroni’s multiple comparisons test (two sided). **(C)** Schematic showing behavioral setup with hindlimbs placed on a vibrating platform. During, and after the light exposure with implanted miniature-LEDs, their sensitivity to the vibrotactile stimulation was tested again. Bottom, structure of the 2AFC behavioral task. During stimulus trials, mice licking right following stimulus onset were rewarded and classified as hits. During no stimulus trials, licks right were recorded as miss. d’ were calculated to assess performance. **(D)** Top, v-shaped perceptual sensitivity curves before and during the optogenetic inhibition. Amplitude thresholds as a function of vibration frequency for mice (shaded lines: individual mice, symbols: mean). Bottom, the behavioral effect of PC-LSCs inhibition at individual frequencies. One-way ANOVA, P < 0.05, Bonferroni’s multiple comparisons test (two sided).

Next, we examined the role of ETV+ LSCs within the Pacinian corpuscle in regulating the perceptual thresholds of mice in a vibrotactile detection task. We adapted a tactile perception task for water-restricted, head-restrained mice (Prsa et al. 2021) in which ETV-ArchT mice were trained to report a high frequency vibration stimulus delivered to the hindpaw (**Fig. 5C**, see **Methods** for details). Following 12-25 days of behavioral training mice could correctly report detection of the stimulus by licking a water spout within a 2-s response time window (**Fig. S8**). After training, we implanted miniature-LEDs into the fibula region of both hindlimbs, controlled remotely through a custom made wireless optogenetic device (BlueBerry, see **Methods** for details), in order to inhibit ETV-ArchT+ LSCs in PCs during behavioral tasks. After a few days of recovery, we retested the mice in the perceptual task during light exposure. The detection tasks yielded comprehensive sensitivity curves and mice with LSCs inhibited by exposure to yellow light showed a reduction by a factor of five in their ability to correctly detect the stimulus across all frequencies (**Fig. 5D**). This effect was reversible, as post-exposure tests showed that the mice had recovered their perceptual performance back to control levels (**Fig. 5D**). To test whether the effects of yellow light itself, in the absence of ArchT, could activate LSCs, we trained an additional mouse that lacked ArchT expression (**Fig. 5D**, lower panel). Consistently, the mouse showed no changes to its perceptual threshold following an identical procedure of yellow light exposure of the hindlimbs. Together, these data demonstrate that LSCs in Pacinian corpuscles are essential to achieve the lowest threshold levels for high frequency vibrotactile stimuli.

## Discussion

Pacinian corpuscles are essential for the detection of various types of external as well as internally generated stimuli (Birznieks and Vickery 2017; Bell, Bolanowski, and Holmes 1994; Turecek and Ginty 2024). We reveal their intricate, convoluted, multilayered and connected morphology. Reconstructions of multiple cells show how their layers intertwine, and wrap around the axon in a hair-claw like manner, which is similar to the inner cores of avian Pacinian corpuscles in the accompanying study (Ziolkowski, Nikolaev et al., 2024). In both cases, the afferent contains dense protrusions which extend through the cleft structure. A comprehensive understanding of the ultrastructural features of LSCs is not only crucial for refining our understanding of the physical properties of the PC, but will also guide future experiments to study the developmental stages of PCs. Currently, very little is known about how PCs develop and how their function evolved. We also explored the relationship between the positions of the LSC nuclei and the spatial distribution of the LSC-axon contacts (**Fig. 2E-H**) and, surprisingly, revealed a strong spatial correlation between the LSCs’ nucleus position and their first contact to the axon **(Fig. 2I)**, which suggests a “first-come first-served” characteristic, potentially related to developmental mechanisms of the inner core **(Fig. 2I)**. It is possible that the LSCs closer to the ultraterminal developed later than the LSCs closer to the preterminal **(Fig. 1A)**, and the later-developed LSCs can only make contact with the axon in the distal region. This is consistent with the observation that the location of each LSCs passage through the cleft also follows each LSC’s nucleus position **(Fig. 1F)**.

Other than making contact with the axon, a major feature of LSCs is the densely packed thin lamellae (53 layers on average). Our observation that LSC-LSC contacts can be predicted by the surface area of two LSCs **(Fig. S5)**, suggests that direct communication (gap-junctions) between LSCs can be linearly increased by enlarging the surface area, resulting in these highly convoluted structures. Our results allow an estimation of ∼69 gap junctions and ∼97 desmosomes per LSC (totaling 4142.5 gap junctions and 5805.1 desmosomes in the whole inner core), suggesting that LSCs appear to maximize their contact areas with each other for rapid high-fidelity functional connections, but the lack of gap junctions between LSCs and the axon suggest prioritization of mechanical interactions with the axon. This is well-supported by the accompanying study (Ziolkowski, Nikolaev et al.), which demonstrates that injection of the gap-junction permeable dye, lucifer yellow, into inner core LSCs permeates all of the inner core lamellar cells, but does not permeate into the axon. Therefore densely packed lamellae might not only facilitate the transmission of physical vibration (Bell, Bolanowski, and Holmes 1994; Biswas, Manivannan, and Srinivasan 2015) but, at the same time, enhance the information exchange within these complex networks of LSCs, for rapid, high-fidelity, and synchronous communication.

Our discoveries from electrophysiological experiments suggest that LSCs play a pivotal and direct role in neural coding of Pacinian corpuscles. Nevertheless, the lamellated architecture of Pacinian LSCs presented in this, as well as in the accompanying study (Ziolkowski et al., 2024), leads to the question: by what mechanisms do the complex LSC structures influence the physiological responses of PCs? Although we observed clear physiological impacts of optogenetic manipulation in LSCs on the coding of vibration in PCs (**Fig. 4**), future *in vivo* experiments will be required to visualize membrane activity and physical movements during vibrotactile and optogenetic stimulation. Such experiments could greatly advance our understanding of how LSCs participate in the complex process of mechanotransduction and the exquisite vibration frequency tuning of PCs.

Finally, we demonstrated reduction in the neuronal vibration detection thresholds upon optogenetic activation of LSCs, which is partially consistent with the accompanying study which showed electrical activation of LSCs decreases the threshold of mechanical activation of the avian Pacinian afferent (Ziolkowski et al., 2024). Additionally, we demonstrate the elevation of vibrotactile perceptual thresholds by a factor of 5 upon optogenetic inhibition of hindlimb LSCs (**Fig. 5**), suggesting that LSCs in PCs are essential for detection of vibrotactile stimuli at the lowest perceivable vibration amplitudes. Overall, our results establish LSCs as a critical player in the exquisite sensitivity to high-frequency vibrotactile stimuli, facilitated by PCs (Ziolkowski et al., 2024). In future, it will be interesting to investigate if higher vibrotactile sensitivity thresholds could impact gait, locomotion, dextrous motor control and bodily self-awareness (Turecek and Ginty 2024).

## Methods

### Animals

All electron microscopy experiments were carried out in double-transgenic mice. Homozygote Ai14 males carrying a floxed tdTomato fusion gene inserted in the Gt(ROSA)26Sor locus in a C57BL/6 strain (Madisen et al. 2010) (Jackson Laboratory, stock no. 007914) were mated with heterozygote ER81/Etv1-CreER females expressing the CreERT2 fusion protein from the ER81/Etv1 promoter elements (Taniguchi et al. 2011) (Jackson Laboratory, stock no. 013048). The Cre-mediated recombination resulted in the expression of the floxed tdTomato sequence in the ER81/Etv1-expressing cells of the offspring. Amongst other cells, ER81/Etv1 is expressed in the inner core region of Pacinian corpuscles (Fleming et al. 2016). To induce CreER-based recombination, double-transgenic adult offspring were administered five consecutive daily doses of 100 μL tamoxifen (T5648, Sigma-Aldrich) solution (20 mg/mL) dissolved in corn oil (C8267, Sigma-Aldrich) by peritoneal injection. Electrophysiology experiments on nerve afferents were conducted with C57BL/6 (Charles River Laboratory) mice. Wild type mice were purchased from Charles River and transgenic mice were obtained from The Jackson Laboratory and bred in the animal facility of the University of Geneva. Experiments involving optogenetic manipulation of Schwann cells were conducted in double-transgenic mice generated by mating homozygous Ai32 mice carrying a floxed *ChR2*-EYFP fusion gene inserted in the *Gt(ROSA)26Sor* locus in a C57BL/6 strain (Jackson Laboratory; stock no. 012569) or homozygous Ai40 mice carrying a floxed *ArchT*-EYFP fusion gene inserted in the *Gt(ROSA)26Sor* locus in a C57BL/6 strain (Jackson Laboratory; stock no. 021188) with heterozygote ER81/Etv1-CreER mice.

The mice were housed in an animal facility in groups of maximum 5 animals per cage and maintained on a 12:12 light/dark cycle. All experiments were performed during the light phase of the cycle. The animals did not undergo any previous surgery, drug administration or experiment. All procedures were approved by the Institutional Animal Care and Use Committee of the University of Geneva and Geneva veterinary offices.

### Surgery

15 to 30 week old mice were surgically prepared for terminal electrophysiology. Surgeries were conducted under isoflurane anesthesia (1.5 to 2%) and additional analgesic (0.1 mg/kg buprenorphine intramuscular (i.m.)), local anesthetic (75 µL 1% lidocaine subcutaneous (s.c.) under the location for incision) and anti-inflammatory drugs (2.5 mg/kg dexamethasone i.m. and 5 mg/kg carprofen s.c.) were administered. Mice were fixed on a bite bar with a snout clamp and rested on top of a heating pad. All electrophysiology experiments were surgically prepared under terminal anesthesia by inhalation of isoflurane (∼2%) and the body temperature was maintained near 37 °C. For analgesia, buprenorphine (0.1mg/kg SC) was provided 15-20 min before the procedure, except for behavioral experiments, where chronic preparation was employed. An additional dose of buprenorphine was given if the procedure exceeded 4 hours. The whole procedure was limited to 8 hours. For chronic optogenetic experiments, a custom-made titanium head bar and a small headstage for wireless multichannel optogenetic system were fixed on the skull with a cyanoacrylate adhesive (ergo 5011, IBZ Industrie) and dental cement to allow subsequent head fixation. After exposing the PCs with a small incision, miniature-LEDs were sutured to the tissue right on top of the fibula bilaterally, and were connected to the headstage with insulated bronze cables threaded underneath the skins (Kathe et al. 2022). The incision then was closed by suturing. After the initial surgical phase, the animal was maintained under light isoflurane anesthesia (0.25 – 0.75%) and allowed at least 2 days of recovery before recording.

### Electrophysiology

For electrophysiology experiments, mice were anesthetized by isoflurane inhalation (2%) and body temperature was maintained near 37°C with a heating mat. For analgesia, buprenorphine (0.1mg/kg SC) was provided 15-20 min before the procedure. To assess the sensory afferents innervating Pacinian corpuscles, a skin opening was performed along the hamstring muscle. Then, the sciatic nerve was carefully isolated using ophthalmic scissors and tweezers. The pia mater spinalis and dura mater around the nerve were removed, and finally, the nerve bundle was separated into single fibers (15-20 μm in diameter) and placed on two Ag/AgCl wire electrodes in the recording pool filled with mineral oil. A third ground Ag/AgCl electrode was placed in the muscle next to the recording chamber.

To characterize the response properties of sensory afferents at the peripheral level, we recorded stimulus-evoked activity of afferents *in vivo* using hook electrodes in the tibial nerve. Vibratory stimuli were applied to different locations of the hindlimb or forelimb using a calibrated piezo stimulator (Physik Instrumente P-841, E509 controller, E504 amplifier), so that the neural responses to all combinations of locations, frequencies and amplitudes could be characterized. To determine the mechanical sensitivity threshold and frequency tuning of each afferent, a series of sinusoidal mechanical stimuli (duration 20 s for frequencies above 100 Hz; linearly increasing amplitude) were applied to the hand-mapped receptive field of individual afferents with the piezo actuator. In order to study purely vibration induced responses, the probe was first placed on the skin for a period of time of a few seconds to minutes and the stimulation for threshold mapping was slowly increased.

The signal was amplified and filtered (>10 Hz and <10 kHz) and acquired at 30 kHz (PXIe-1073, National Instruments) using WaveSurfer (https://wavesurfer.janelia.org/) Matlab (Mathworks) routines. Trial start, stimulus onset triggers, and details of stimulus were saved in parallel on separate channels and used for *post-hoc* alignment of recorded spikes. After the recording, animals were euthanized by overdose with isoflurane (5%) followed by cervical dislocation and bleeding.

### Vibrotactile stimuli

The vibrotactile stimuli were generated by a bimorph piezoelectric multilayer bender actuator with a piezoelectric stack actuator (P-841.3, Physik Instrumente) equipped with a strain gauge feedback sensor for high frequency stimulations (above 100 Hz). A hand-cut blunt plastic cone was mounted on the actuator. The actuator and sensor controllers (E-662, E-618.1 and E-509.S1, Physik Instrumente) operated in an open loop with a contacting force below 50 mN on the mouse skin. The forces of our vibrotactile stimuli measured at the static states of 3µm, and 30 µm, were less than 10 mN, and 10-30 mN. Pure sinusoids (250 or 500 ms duration) of a wide range of frequencies (100 to 1900 Hz) calibrated to produce a desired displacement amplitude (0.01 to 20 µm) were sampled at 10 kHz (USB-6353, National Instruments) and fed to the actuator controller. The sensor measurements were continuously acquired and recalibration of motor commands was regularly performed for the stimuli to remain highly consistent, by comparing the ground truth data acquired optically with a laser doppler vibrometer system (Polytec, OFV-5000). The spectrum of the acquired sensor measurements was analyzed to ensure the integrity of their frequency content (Prsa et al. 2019; Lee et al. 2023).

### Optogenetic stimuli

Optogenetic stimulation of Schwann cells in Pacinian corpuscles (PC-SCs) was generated by blue light illumination (473 nm, 50 mW; Obis LX FP 473, Coherent) out of a fiber (3.5 µm core diameter, 0.045 NA) operated in analog control mode or by yellow light illumination (568 nm, 500 mW; Sapphire LP/LPX, Coherent) fed into an optic fiber with achromatic FiberPorts (Thorlabs), gated with a shutter. The laser power or the shutter was controlled by analog signals from the WaveSurfer (https://wavesurfer.janelia.org/). For the blue laser, the stimulus was a continuous square pulses (2000 ms duration). The mean power of the laser for continuous stimulation used in our experiments was 0.3 mW for blue and 0.5 mW for yellow (measured at the tip of the fiber).

We used a custom-made wireless multichannel optogenetic system (BlueBerry) to apply the stimulation through miniature-LEDs (0402 SMD-QBLP595-IG) in the fibula region. The BlueBerry is made of a low-power Bluetooth module (RN4871-microchip) that communicates the stimulation protocols (frequency, duty cycle, LED channel) with a microprocessor (ATTINY 85) connected to both miniature-LEDs. A mobile application (BLE Terminal) is used to transmit all the stimulation parameters to the bluetooth module of the BlueBerry. For optical stimulation of Pacinian corpuscles, the stimulus was 3 ms short square pulses flashing in 20 Hz with approximately 3 mW.

### Serial block face scanning electron microscopy (SBF-SEM) and transmission electron microscopy (TEM): Euthanisia, Perfusion and Dissection

Adult mice (15 - 30 weeks) were anesthetized with sodium pentobarbital (60 mg/kg) by intraperitoneal injection. They were subsequently transcardially perfused with 15 ml of 0.1M sodium phosphate buffer, followed by 60 ml of 2% paraformaldehyde (15710-S, EMS) 1% glutaraldehyde (G7526, Sigma-Aldrich) in 0.1M sodium phosphate buffer pH 7.4. The perfused animals were placed in a sealed plastic bag for 1-2 hours before the dissection of Pacinian corpuscles was carried out. Dissection of Pacinian corpuscles (**Fig. S1**) was carried out under an fluorescent stereo microscope (Leica M165FC). First, the four limbs were isolated by cutting above the elbow/knee and the skin was cut away until the wrist/ankle. The paw was pinned onto a dissection tray with the palm/sole facing up, and then muscles and tendons were gently cut away to reveal the ulna or fibula and tibia. To identify the PCs, the expression pattern of tdTomato was revealed by illuminating the bone green LED light and using an ET TxRed filter set (Leica). The PCs present in bundles (resembling a cluster of grapes), and could be dissected from the periosteum by gently lifting them off the bone with fine forceps (Dumont #5SF Forceps, FST) and pulling the bundle up where the afferents converge. Once dissected, the PCs were post-fixed overnight at 4°C, by immersion in 2% paraformaldehyde (15710-S, EMS) 1% glutaraldehyde (G7526, Sigma-Aldrich) in 0.1M sodium phosphate buffer pH 7.4.

### SBF-SEM: Sample processing and imaging

For SBF-SEM, the sections were post-fixed, stained and embedded according to a protocol similar to Hua et al. (2015). This was done by the BioEM Facility of the Ecole Polytechnique Federale Lausanne (EPFL) (Hua, Laserstein, and Helmstaedter 2015). In brief, the samples were first post-fixed in 1.5% potassium ferrocyanide and 2% osmium in 0.13M ice cold cacodylate buffer. Then they were stained sequentially by 1% thiocarbohydrazide (40 minutes at 40°C), 2% osmium tetroxide (90 minutes at room temperature), 1% uranyl acetate (overnight at 4°C) and finally lead aspartate solution (120 minutes at 50°C). This was followed by dehydration in increasing concentrations of ethanol (50%, 70%, 96% x 2, 100% x 2) and then embedding in Durcupan resin. The samples were left to harden at (65°C).

Sample quality was confirmed and regions of interest were identified using transmission electron microscopy (TEM). The blocks were then trimmed and glued to an aluminum stub using conductive resin. The imaging was carried out in a scanning electron microscope (Zeiss Merlin, Zeiss NTS) containing an ultramicrotome (3View, Gatan), also at the BioEM Facility of the EPFL. Two PCs were imaged. For both, an acceleration voltage of 1.7 kV and dwell time of 1 µs were used. An acceleration voltage of the 1.7 kV was used with a pixel size of 6.5 nm with a dwell time of 1 µs. Two PCs were imaged. For the first (“PC1”), a pixel size of 40 nm was used, and the resin slice cut between each image had a thickness of 200 nm. The total size of the block was 138 x 145 x 273 µm. It contained a full PC. For the second PC (“PC2”), a pixel size of 7 nm was used, and the resin slice cut between each image had a thickness of 50 nm. The total size of the block was 29 x 31 x 80 µm.

### SBF-SEM: Image processing

Initially image stitching and alignment was done in the TrakEM extension of FIJI imaging software. For “PC1” two tiles were acquired, these were stitched using the “Montage multiple layers” feature of TrakEM. Then, the “Align layers” feature was used to align all layers, for both “PC1” and “PC2”. Where necessary, small manual adjustments were made. These images were then exported into the software Amira (Thermo Fisher Scientific), which was used for segmentation. For “PC1”, the axon, inner core region, nuclei of inner core cells, intermediate layer region, 14 outer core layers, capsule region and myelin sheath were reconstructed (**Fig. S2**). For “PC2”, the axon (**Fig. S2**), as well as an inner core cell (**Fig. S3**), were reconstructed in a 25 µm sub-segment . The segments were visualized in Amira using the “Generate Surface” tool.

### SBF-SEM: Machine learning assisted image segmentation

To extract individual LSCs in this electron microscopy dataset, the stacks of electron microscopy images were first processed using automated segmentations via Webknossos automated segmentation services, followed by manual proofreading. Initially, a Convolutional Neural Network (CNN) was trained using our manual segmentations. Subsequently, the trained model was employed to generate affinity predictions, indicating the likelihood of a voxel being connected to its neighboring segments. Volume segments were then grouped into agglomerates ranked based on their predicted affinity (Funke et al. 2019) with additional restraints to reduce the occurrence of merge errors (e.g. each segment contained only one nucleus). Furthermore, we conducted human proofreading to the fullest extent possible to rectify merge or split errors before proceeding with the data analysis of these annotations.

### Transmission Electron Microscopy (TEM): Sample processing and imaging

After overnight post-fixation, the samples were prepared for TEM as follows. Each sample was placed in a 10 mL glass test tube with a cork lid for the whole protocol. The samples were washed at room temperature in 0.1M PB pH 7.4. Then, they were post-fixed in 2% osmium in the Millonig buffer for 1 hour at 4°C, followed by washing at room temperature with ddH_2_O. The samples were then stained overnight with 0.25% uranyl acetate at 4°C. The next morning, the samples were washed with ddH_2_O. They were then dehydrated in ethanol (15058, Reactolab SA), increasing concentration from 30%, 50%, 70%, 90% to finally 100%. This was followed by 100% propylene oxide (82320, Fluka Analytical), twice for 10 minutes. In the mean time, the epoxy resin was prepared. For 100g, the following were mixed: 46.34 g Epoxy embedding material (45345-250ML-F, Sigma Aldrich), 27.76 g DDSA (45346-250ML-F, Sigma Aldrich), 25.9 g hardener MNA (45347-250ML-F, Sigma Aldrich) and 1.5 g Accelerator (45348-250ML-F, Sigma Aldrich). The samples were then left in a 50% propylene oxide and 50% epoxy resin mixture for 30 minutes and then left in 100% epoxy resin mixture overnight, both at room temperature. Finally the samples were moved from the glass test tubes into a silicone mould with fresh 100% epoxy resin mixture. The samples were left to cure for 72 hours at 60°C.

The resulting resin blocks were trimmed and ultrathin sections were cut at a thickness of ∼60nm using a diamond knife (DiATOME Ultra 45°) and ultramicrotome (Leica Ultracut UCT).The resin blocks were mounted in the ultramicrotome so that the Pacinian corpuscles would be cut along the coronal axis. The cut slices were collected on slot grids that had been prepared at the Electron Microscopy Facility of the University of Lausanne. The sections were post-stained with 5% uranyl acetate for 10 minutes and lead nitrate for 7 minutes.

Imaging was carried out on an FEI Morgagni transmission electron microscope. For an overview of the full Pacinian corpuscle, images were taken at a magnification of 2800x. To image the inner core, images were taken at a magnification of 8900 to 14 000x. For higher resolution of details, images were taken at a magnification of 28 000x. When the region of interest was larger than the field of view, multiple tiles were taken to cover the whole area of interest.

### TEM: Image processing

Images were cropped and adjusted for optimal contrast using Fiji/ImageJ. If multiple tiles were taken, the Plan Brightness Adjustment plugin of Fiji was applied to all images, then they were aligned and stitched together in the TrakEM extension of Fiji. The TEM images presented in this manuscript were pseudo-coloured by hand using Illustrator.

### Cryo temperature staining and embedding for TEM

The perfused, chemically fixed samples were cryo-protected in a solution of 2% glycerol and 20% DMSO, in 0.01M PBS, and then high pressure frozen in the same solution (Leica MicroSystems ICE high pressure freezer). They were then placed in plastic cryo tubes inside a low temperature embedding machine (AFS, Leica Microsystems) held at -90°C. These tubes contained acetone with 2% osmium tetroxide, 0.2% uranyl acetate, and 5% water. The temperature was then raised to -20°C over three days, and then the solution was exchanged for pure acetone while the temperature was raised again to 0°C. At this point the acetone was exchanged for a 50/50 mixture of epon resin and acetone, and the temperature then brought up to 20°C. After several exchanges of pure resin the samples were then flat embedded between glass slides and the resin hardened for 24 hours at 65°C.

### Detection task in mice

In order to determine their perceptual thresholds, we used a two-alternative forced choice task to train seven mice in a vibrotactile detection task (Prsa et al. 2021). Mice were trained to lick, in the response period, toward either a right or left reward spout if a vibrotactile stimulus was present or absent during the preceding stimulus period, respectively. All other experimental conditions were as described in the previous study (Prsa et al. 2021). Correct responses were rewarded with a drop of water at the corresponding spout and incorrect responses were not punished by a timeout. Trials without a response were neither rewarded nor punished and occurred on <5% of trials. To minimize a direction bias, the trial type was chosen pseudo-randomly by allowing a maximum of three trials of the same type in a row (50% chance of occurrence for each otherwise). We tested the perceptual thresholds at seven different frequencies (300, 450, 700, 1000, 1300, 1600, and 2000 Hz) in separate sessions and in randomized order. Between 1 and 3 sessions were tested in a single day and the same session (i.e., frequency) was repeated up to five times on separate days per mouse. Prior to testing, the mice were first trained on all frequencies at the largest possible amplitude that the actuator could produce at each frequency. This value ranged from 10 µm (at 300 Hz) to 1 µm (at 2000 Hz). The training lasted 12–24 days. Testing of each frequency started at the largest possible amplitude and was progressively attenuated in −4 dB steps after every six vibration trials (total of ≈12 trials) if the proportion of correct responses exceeded 70%. The amplitude was increased by 4 dB if the proportion of correct responses decreased below 60% after every ≈12 trials (including at least six vibration trials). To determine the perceptual threshold at each frequency, we compared the ratio of correct responses for each bout of trials at a given amplitude to chance (i.e., 0.5) using the one-sided binomial test. The threshold was the lowest amplitude of the session for which the test yielded a significance level of <0.05. The thresholds of repeated sessions were averaged and allowed establishing the V-shaped vibrotactile sensitivity curves.

### Data analysis

#### Significant responses

For electrophysiological recording, tuning curves were computed by the spike rate of the neurons. To ensure that we were recording from a single nerve fiber/neuron and to avoid the complications of spike sorting, we only sampled units with a signal-to-noise ratio (SNR) > 5, when responding to a vibration stimulation (100 or 700 Hz). SNRs were calculated as the maximum amplitude of the mean spike waveform divided by the standard deviation of the background noise. There was no additional offline filtering performed for spike detection.

#### Tuning curve fitting

Neuronal structures were considered to be responsive if the maximum stimulus-related spiking responses to any stimulus was greater than 5% on average, and also greater than 2 standard deviations (SD) above the mean firing rate. In addition, we required that neurons responded at least 2 SD above baseline on at least 20% of the trials tested. The same criteria were used to identify neurons with significant responses to the vibration, optogenetic and stimuli. Neuronal structures were considered to be frequency selective if they were responsive and also met the following criteria: (1) well fit by the polynomial function (r > 0.7, p < 0.05), and (2) tuning index (*TI*) > 0.2

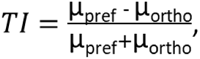

where μ_pref_ denotes the mean response to the preferred stimulus frequency and μ_ortho_ is the mean response to the least preferred stimulus frequency, defined by the curve fitting. To characterize a neuron’s tuning to vibration frequency, the normalized mean responses were fit to frequency by the polynomial curve fitting function (Matlab) using the method of non-linear least squares, with the degree of polynomial fit set at 6. The tuning preference of individual neurons was defined by the peak of the curve and the tuning width was defined by the half-width at the half-maximum of the curve (HWHM).

#### Phase-locked spiking

To determine whether a nerve fiber or a neuron was entrained by the sinusoidal vibration, we calculated standardized inter-spike intervals. We chose this measure because it is immune to variability in response onset times across different stimulus repetitions. For a given vibration frequency, the inter-spike intervals (ISI) of all possible spike pairs that occurred during stimulation were calculated and grouped across stimulus repetitions. Because entrained spiking should yield an ISI distribution that peaks at integer multiples of the sinusoidal stimulus period *T*, values were converted to standardized inter-spike intervals (*S*ISI) according to *S*ISI = *T* + (ISI − *nT*), where *nT* is the integer multiple of *T* closest to ISI. Entrainment probability was defined as the percentage of ISIs in the [*nT* − *T*/12, *nT* + *T*/12] interval. Given that standardized ISIs are distributed between *nT* − *T*/2 and *nT* + *T*/2, entrainment probability should equal unity in the case of perfect entrainment and be equal to 1/6 in the case of chance entrainment. For each neuron and at each frequency we repeated the calculation of entrainment probability 1,999 times with randomly sampled ISIs (with replacement). We then measured whether the lower 99^th^ percentile of the repeated measures was less than 1/6, which constitutes a one-tailed bootstrap test at significance level P < 0.01.

#### Quantification of surface areas of individual lamellar Schwann cells and the axon

To quantify the surface areas of individual lamellar Schwann cells and the axon, all annotations were first downloaded through WebKnossos as TIFF stacks. These were followed by the extraction of individual segmentations using Fiji/ImageJ, based on their “Segment ID” labeled in WebKnossos, separately. After setting the pixel-to-micron scale, the ‘‘Analyze Particles’’ function of Fiji/ImageJ was applied to process all EM images and export the perimeter for the selected segment (Schwann cell or the axon). The surface area for that selected segment was calculated by multiplying the total perimeter by the z-step size of 0.05 µm (Surface area (µm²) = total perimeter (µm) * 0.05 (µm)). To quantify the surface areas in different quadrants (transverse view) or along the longitudinal direction, the annotations were cropped or selected before processing with the ‘‘Analyze Particles’’ function.

#### Quantification of LSC-Axon and LSC-LSC Contacts

LSC-axon contacts occur at the axon protrusion or body. Observations indicate that lamellar Schwann cells (LSCs) and the axon protrusion engage in light, transient "kiss-and-run" interactions, while LSCs contact the axon body in a patchy pattern. We manually determined the first contact position, number of contacts (distance < 30 nm), and contact range between individual LSCs and the axon protrusion in all EM images. The estimated size of each contact is ∼0.1 μm. The total contact area with the axon protrusion was calculated by summing the contact range (μm) and multiplying by 0.1 μm. To determine the total contact area of an individual Schwann cell with the axon (protrusion and body), we used the "MorphoLibJ" plugin in Fiji/ImageJ. We performed morphological dilation to expand the axon boundaries by 30 nm, measured the overlap perimeter between the dilated axon and LSCs, then multiplied half of this perimeter by the z-step size of 0.05 µm. LSC-LSC contacts were measured similarly to LSC-axon contacts. To determine the contact area of one LSC with other LSCs, we used the "MorphoLibJ" plugin in Fiji/ImageJ. We performed morphological dilation to expand the LSC boundaries by 30 nm, measured the overlap perimeter between the dilated LSC and other LSCs, then multiplied half of this perimeter by the z-step size of 0.05 µm.

#### Correlation Analysis

The Pearson correlation coefficients (r) used in this paper to compare two groups were calculated using GraphPad Prism software. The P value of the correlation coefficient is used to determine the significance of the correlation analysis.

### Statistical analysis

Statistical analyses were conducted using Graphpad Prism or Matlab, with *p < 0.05 set as the significance threshold. Detailed descriptions of the statistical tests and sample sizes for each experiment are provided in the figure legends and main text. No statistical methods were used to predetermine sample size and all experimental animals were included in the analysis. The normality assumption was tested with the Kolmogorov-Smirnov test. Non-parametric tests were used when the normality assumption was not met. We used a two sided non-parametric Wilcoxon rank-sum test or Mann Whitney test to compare two groups and the Kruskal–Wallis test to compare multiple groups with post-hoc tests using Dunn’s test, without assumptions of normality or equal variances. Paired-sample t-test was used to compare the effect of optogenetic manipulations. All statistical methods were two-sided.

### Data and code availability

SBF-SEM dataset is publicly available on Webknossos: https://webknossos.org/links/NxewRCaXI6KD2JKP (raw), https://webknossos.org/links/ItaEln7CJqGS51DY (machine learning assisted image segmentation). All data reported in this study and code used for analysis will be shared by the lead contact upon request.

## Acknowledgements

We thank members in Bagriantsev and Gracheva labs (Yale University) for advice and comments on the manuscript; G. Cuenu for help with histological techniques and mice breeding at the early stages; R. Vickery (UNSW) for advice on electrophysiology experiments; P. Ruga Fahy (PFMU, UNIGE) for assistance with TEM; N. Liaudet (Bioimaging Core Facility, UNIGE) for assistance with the manual segmentation pipeline; C. Genoud (EMF, UNIL) for feedback on EM data and all members of Huberlab for discussion and support.

## Author contributions

K.-S.L., D.T.W. and D.H. conceptualized the study and designed experiments. M.-C.C-C., J.B. and G.K. conducted serial block-face scanning electron microscopy. Y.-T.C. and D.T.W. analyzed data on the electron microscopy. K.-S.L. conducted the physiological and behavioral experiments and analyzed data, with assistance from A.J.L. and A.N. Y-T.C., K.-S.L., D.T.W., A. J. L. and D.H. wrote the manuscript.

## Data and materials availability

All data needed to evaluate the conclusions in the paper are present in the paper and/or the Supplementary Materials, and will be shared by the lead contact upon request.

## Supplementary Information

**Fig S1:**
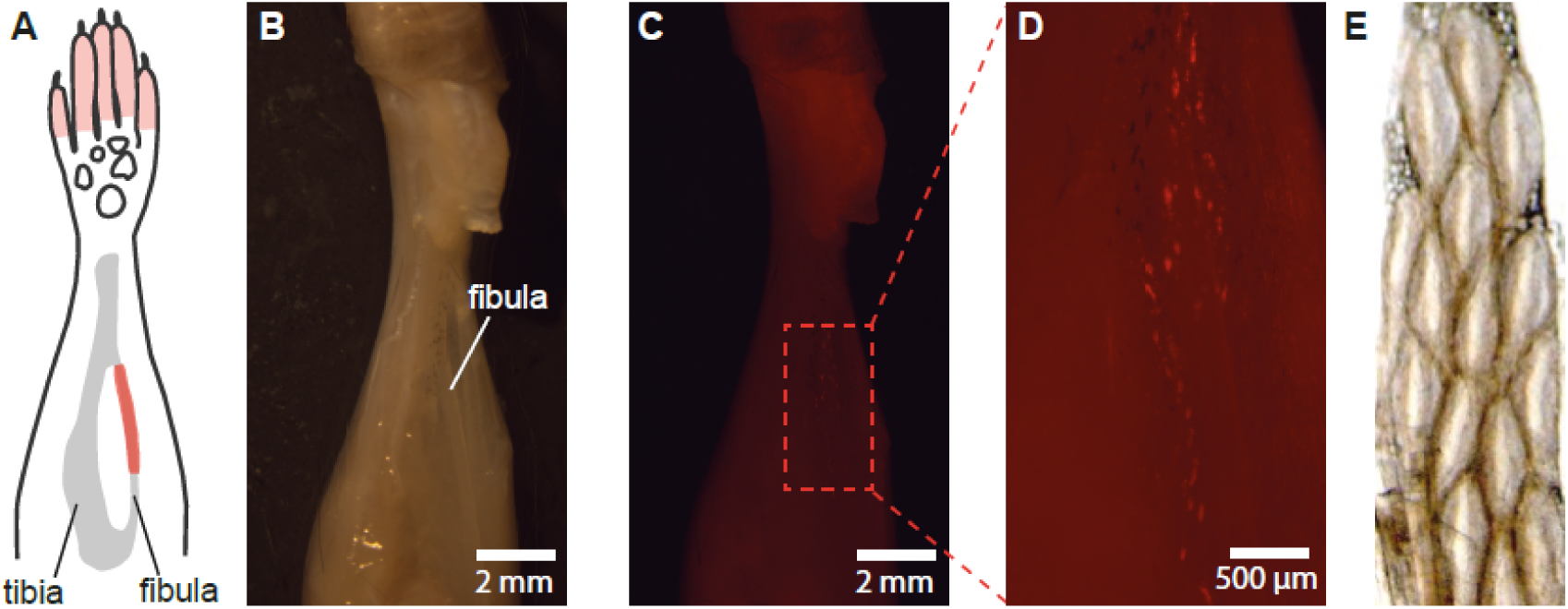
PCs can be reliably identified and dissected from the fibula of the mouse hindlimb. **(A)** Distribution of PCs in the mouse hindlimb. **(B)** Mouse hindlimb with muscles removed to exposed tibia and fibula. **(C-D)** Dissected hindlimb under a TxRed filter, showing the ETV1+ inner core regions of the PCs (each red fluorescent spot is one inner core). **(E)** A bundle of dissected PCs prepared for SBF-SEM, just before resin embedding.

**Fig S2:**
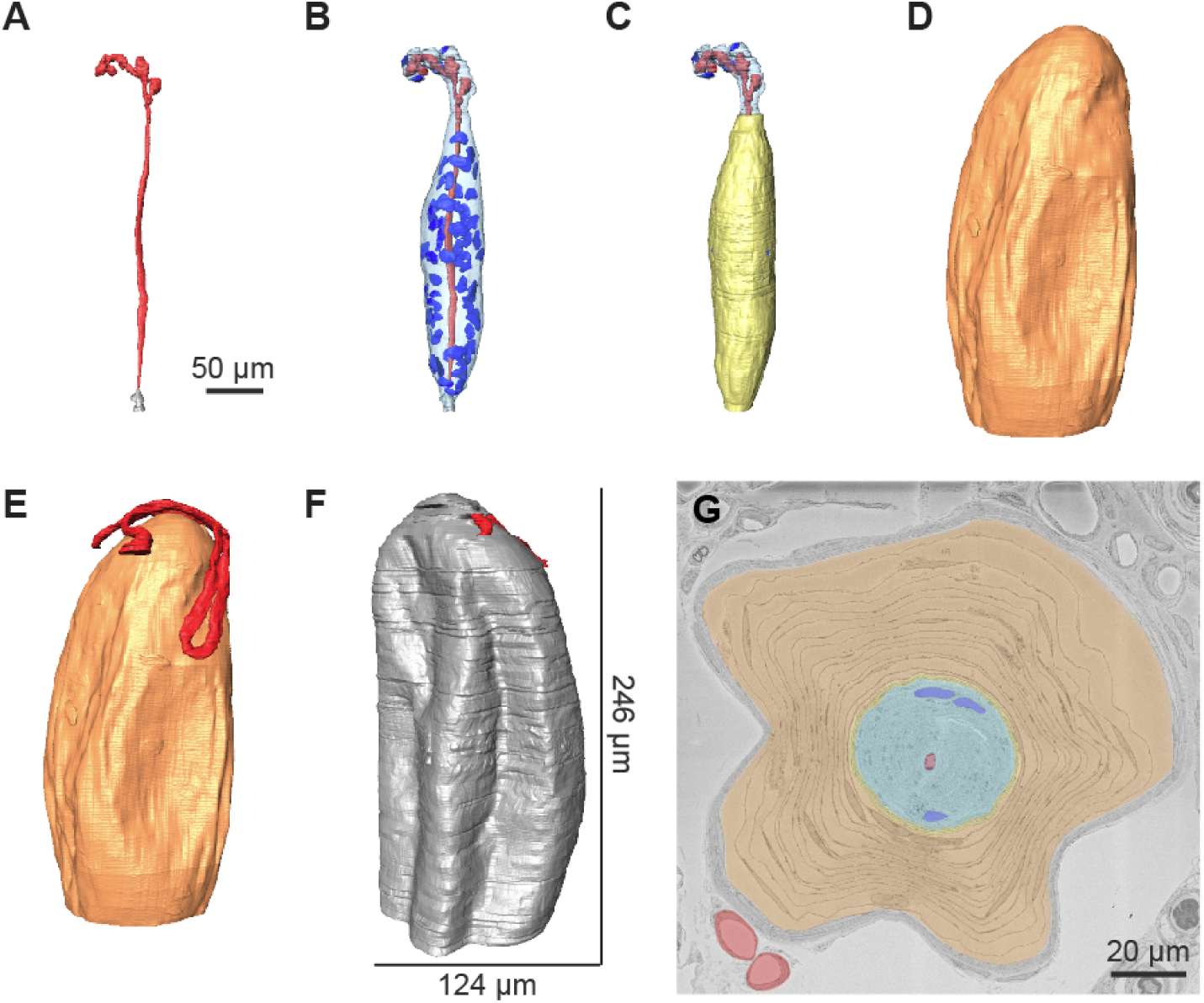
Reconstructions of SBF-SEM data reveal PC structure in 3D. **(A-F)** Manual reconstruction of a full mouse PC, imaged at lower resolution (40 x 40 x 200 nm). The scale bar in B is true for B-G. **(A)** The axon (red) is myelinated (gray) in the pre-terminal, long and oval-shaped in the terminal, and finally bulbing and bifurcating in the ultraterminal. **(B)** The outline of the inner core (transparent blue) region. At this resolution, the nuclei (opaque blue) in the IC could be reconstructed, but not the layers of the LSCs. In this inner core there were 63 nuclei, suggesting it is made up of as many LSCs. **(C)** The intermediate layer (yellow) is uniform in the pre-terminal and terminal, but merged with the outer core layers in the ultraterminal, so that portion could not be reconstructed. **(D)** One of fourteen outer core layers (orange). **(E)** The blood vessel (red) between the outer core and capsule. **(F)** The capsule (gray). At its largest, the PC was 246 μm long and 124 μm wide. **(G)** Pseudo-coloured image from the SBF-SEM stack, showing the different reconstructed areas.

**Fig S3:**
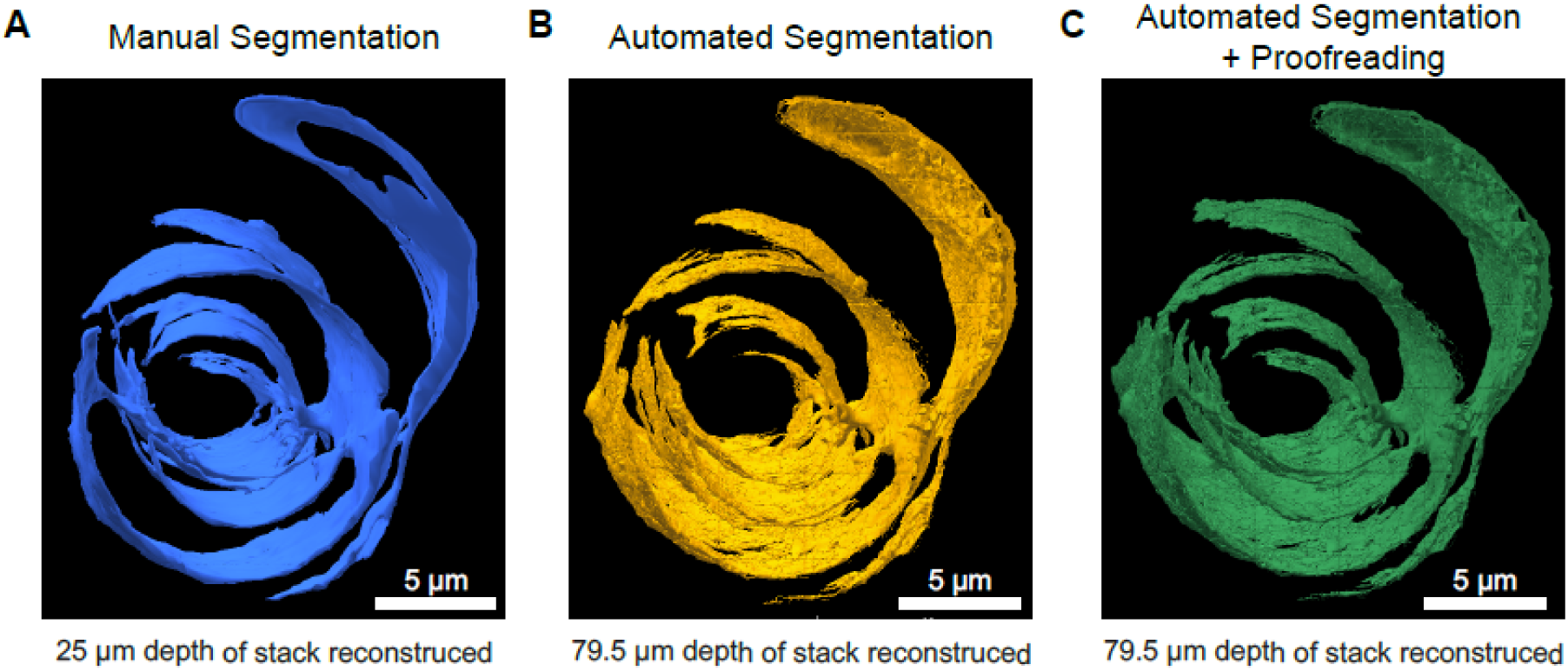
LSCs segmented by automated methods are equivalent to those segmented manually. **(A)** LSC4 when segmented by hand using AMIRA. The segmentation required over 300 hours. **(B)** LSC4 when segmented using the Webknossos automated segmentation services. **(C)** LSC4 when segmented using the Webknossos automated segmentation services, followed by manual proofreading. The manual proofreading required around 10 hours.

**Fig S4:**
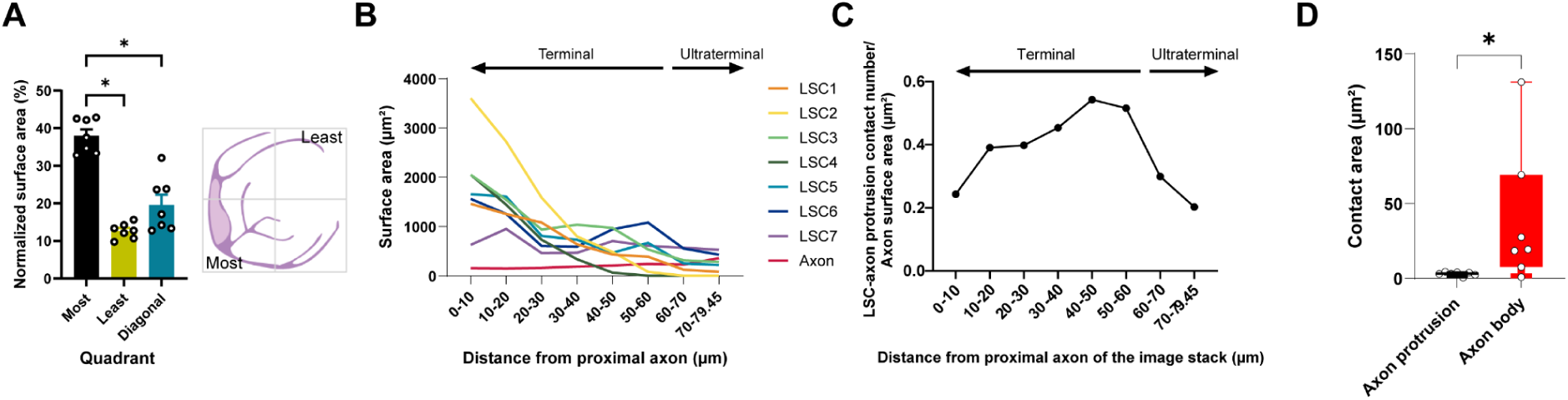
The distribution of seven lamellar Schwann cell surface areas and their axon protrusion contacts. **(A)** The surface area within the quadrant with the most surface area is more than the surface area within its diagonal quadrant and the quadrant with the least surface area. Statistical significance is calculated using T-Tests. *p < 0.05. n= 7. **(B)** The lamellar Schwann cells surface area decreases proximally to distally (statistical significance is calculated using One-way ANOVA. (F(7,48) = 11.88, *P < 0.0001, n = 7), while the axon surface area increases. **(C)** The density of LSC-axon protrusion contacts from proximal to distal. **(D)** LSCs make more contacts with the axon body than with the axon protrusion. Statistical significance is calculated using T-Tests. *p < 0.05. n= 7.

**Fig S5.**
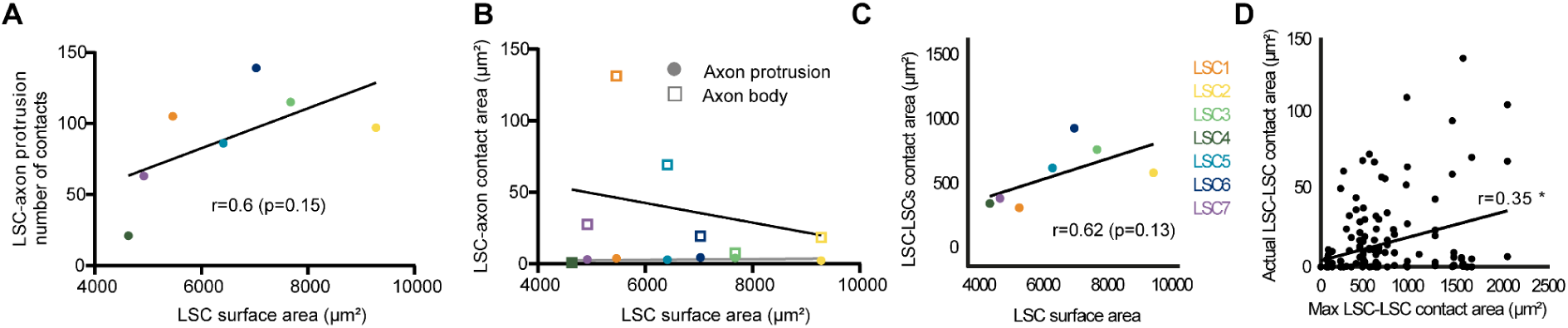
A larger LSC surface area might lead to more axon protrusion contacts but not necessarily to more axon contact area. **(A)** There is a trend of positive correlation between the size of the surface area and the number of axon protrusion contacts. **(B)** There is no apparent correlation between the size of the surface area and the size of the LSC-axon contact area. **(C)** LSC surface area appears to be positively correlated with the size of the LSC-LSC contact area. **(D)** The bigger anatomical presence of two LSCs among the same region can predict the larger LSC-LSC contact area (see **Methods** for details). The relationship between the theoretical maximum LSC-LSC contact area and the measured LSC-LSC contact area is statistically significant.

**Fig S6.**
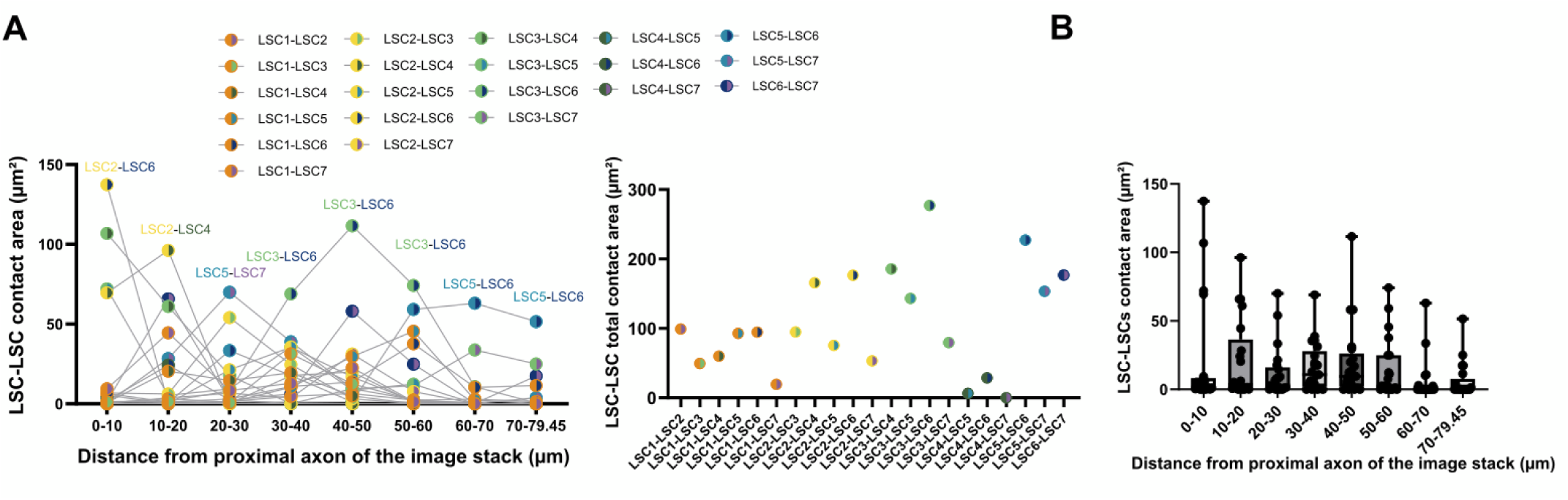
The distribution of LSC-LSC contacts in the PC inner core. **(A)** LSC-LSC contact area measured at different distances from the proximal axon of the image stack, along with their overall contact with the axon. **(B)** The Box plot shows the distribution of LSC-LSC contact areas across the same distance intervals as in **(A)**, and there is no significant difference from proximal to distal (One-way ANOVA; F (7, 160) = 1.279).

**Fig S7.**
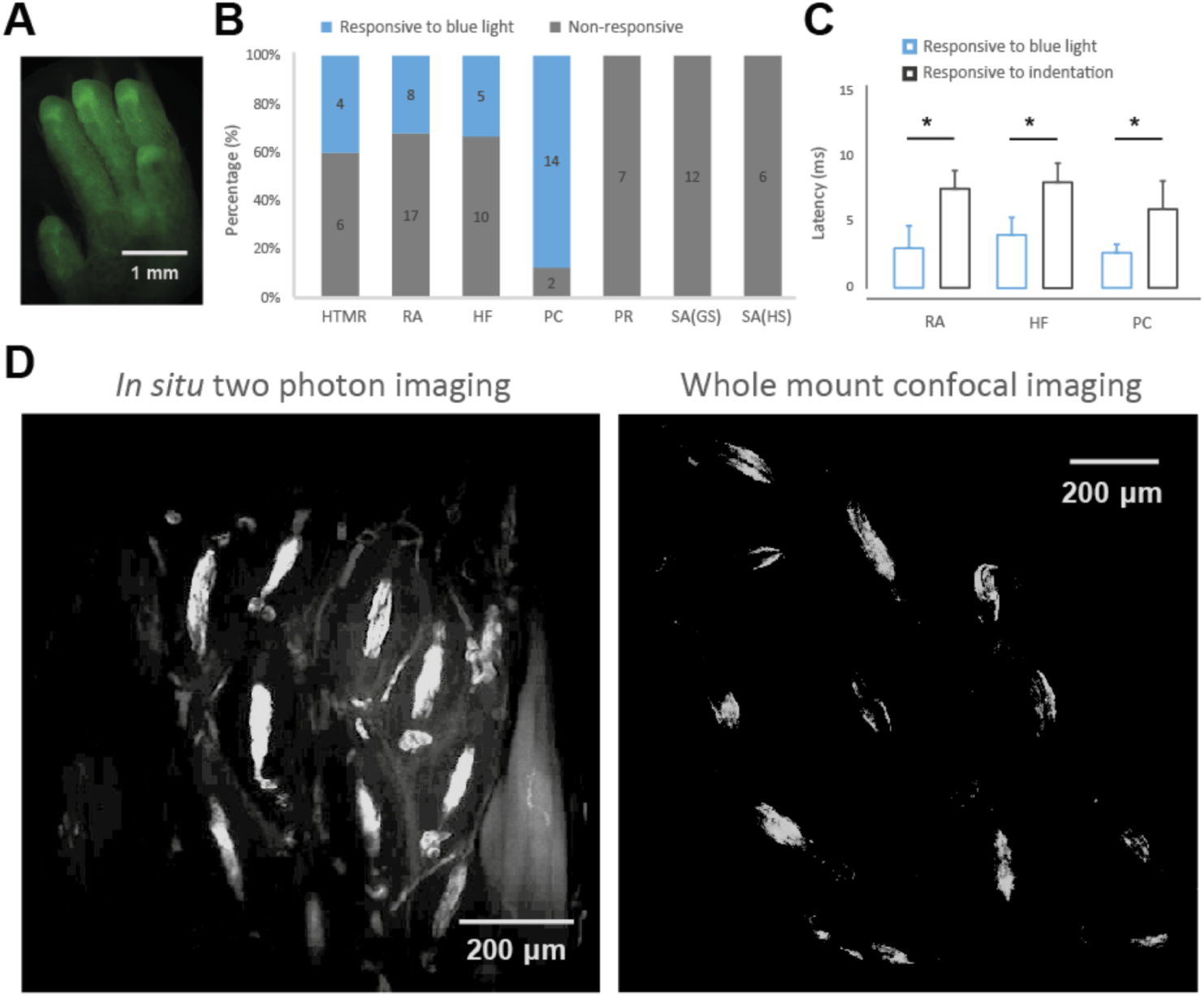
Characterizing the mechanoreceptor types in ETV-ChR2 mice. **(A)** The EYFP signal within the hindpaw digit area. **(B)** Total number of mechanoreceptors recorded in ETV-ChR2 mice, showing proportions of light responsive (blue) and unresponsive mechanoreceptors (gray). HTMR, high-threshold mechanoreceptor; RA, rapidly-adapting receptor; HF, hair follicle; PC, Pacinian corpuscle; PR, proprioceptor; SA, slowly-adapting receptor; GS, glabrous skin; HS, hairy skin. **(C)** First spike latencies from optogenetic activation of Schwann cells and mechanical activation of the same afferent by an indentation (unpaired t-test, *P<0.05). **(D)** Examples of high resolution imaging of intact Pacinian corpuscles from ETV-ChR2 mice showing that EYFP signals are limited to the inner core area, and is absent in the axonal afferents.

**Fig S8.**
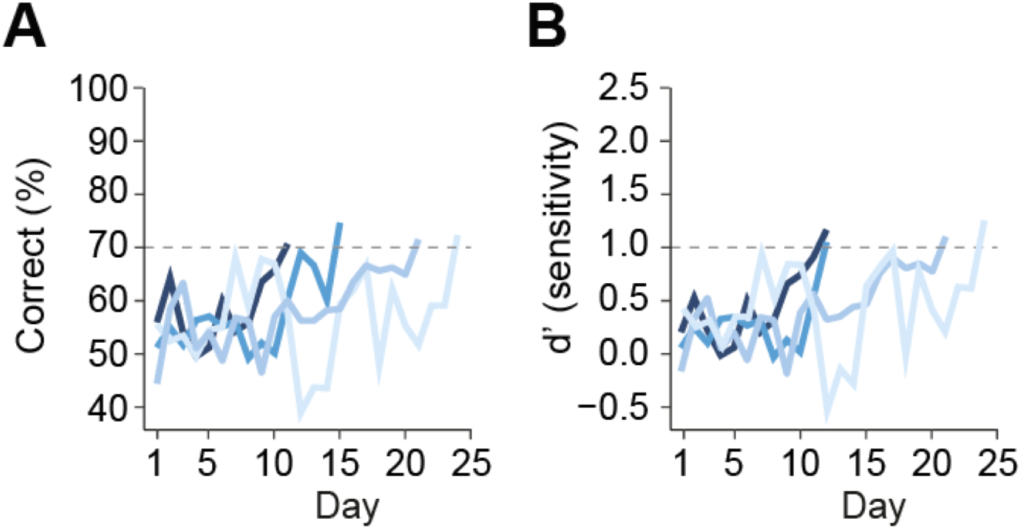
Behavioral training for vibration detection tasks. **(A)** Mice were trained to report the presence of subtle vibrotactile stimuli by licking right and no stimulation by licking left. The correction rate of 5 mice showed a gradual increase over the course of training. **(B)** Mice that reached a performance level of d’ > 1 were considered as successfully trained.

**Supplementary Table 1.**
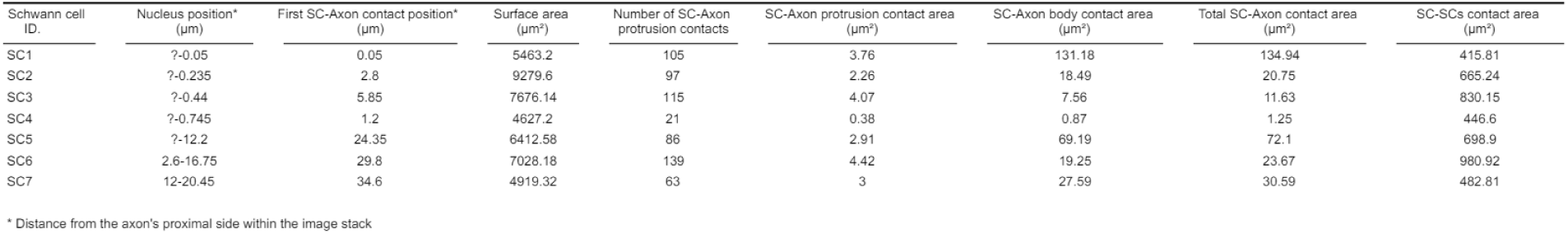
Summary of the structural characteristics of seven Schwann cells.

